# Multiscale predictive modeling robustly improves the accuracy of pseudo-prospective seizure forecasting in drug-resistant epilepsy

**DOI:** 10.1101/2025.09.27.678967

**Authors:** Gagan Acharya, Erin Conrad, Kathryn A. Davis, Erfan Nozari

## Abstract

Extensive research over the past two decades has focused on identifying a preictal period in scalp as well as intracranial EEG (iEEG). This has led to a plethora of seizure prediction and forecasting algorithms which have reached only moderate success on curated and pre-segmented EEG datasets (accuracy/AUC ≳ 0.8). Furthermore, when tested on their ability to pseudo-prospectively predict seizures from *continuous* EEG recordings, all existing algorithms suffer from low sensitivity (large false negatives), high time in warning (large false positives), or both. In this study we provide pilot evidence that *predictive modeling of the dynamics of iEEG features (biomarkers), seizure risk, or both* at the scale of tens of minutes can significantly improve the pseudo-prospective accuracy of almost any state-of-the-art seizure forecasting model. In contrast to the bulk of prior research that has focused on designing better features and classifiers, we start from off-the-shelf features and classifiers and shift the focus to learning how iEEG features (classifier input) and seizure risk (classifier output) evolve over time. Using iEEG from *n* = 5 patients undergoing presurgical evaluation at the Hospital of the University of Pennsylvania and six state-of-the-art baseline models, we first demonstrate that a wide array of iEEG features are highly predictable over time, with over 99% and 35% of studied features, respectively, having *R*^2^ > 0 for 10-second- and 10-minute-ahead prediction (mean *R*^2^ of 0.85 and 0.2). Furthermore, in almost all patients and baseline models, we observe a strong correlation between feature predictability (with some features remaining predictable up to 30 minutes) and classification-based feature importance. As a result, we subsequently demonstrate that adding an autoregressive model that predicts iEEG features on 12 ± 4 minutes into the future is almost universally beneficial, with a mean improvement of 28% in terms of area under pseudo-prospective sensitivity-time in warning curve (PP-AUC). Addition of the second autoregressive predictive model at the level of seizure risk further improved accuracy, with a total mean improvement of 51% in PP-AUC. Our results provide pioneering evidence for the long-term predictability of seizure-relevant iEEG features and the vast utility of time series predictive modeling for improving seizure forecasting using continuous intracranial EEG.

## 1 Introduction

Despite continuous advancements in antiseizure medications, over one-fifth of epilepsy patients remain refractory to pharmacological interventions^1–3^. Motivated by the potential to improve the lives of patients with drug-resistant epilepsy and powered by evidence that epileptic seizures may begin well in advance of clinical onset^4^, seizure prediction and forecasting have been the subject of intensive research efforts over the past two and a half decades^5–12^, all aiming to achieve the same goal: reliably identifying the preictal period with increasing temporal and statistical accuracy. The field has attracted considerable interest, as evidenced by various international initiatives and competitions^13–16^, collectively propelling substantial innovation in algorithmic approaches to seizure prediction. These efforts have resulted in a diverse range of methodologies, from statistical models to deep learning frameworks. Yet, despite this progress, accurate seizure forecasting remains an elusive goal. Existing algorithms universally suffer from low sensitivity, high false positives rates, or both, particularly when tested pseudo-prospectively on con-tinuous EEG. This gap highlights a pressing need for novel approaches that go beyond the commonly-pursued goals of improved feature extraction and advanced classification techniques.

Existing seizure detection, prediction, and forecasting algorithms commonly operate by splitting the time series into (possibly overlapping) segments, extracting certain features (biomarkers) from each, and training a classifier on those features^5–12, 17–20^. The features may be purely signal-based^20–25^ or based on dynamical systems theory^26–33^; they may be derived from time domain^24, 34–36^, frequency domain^34, 37–40^, or both^21, 23, 25, 41,42^; and the classification may be done via a simple threshold crossing^43, 44^ or various state-of-the-art machine learning algorithms^23, 24, 41, 45–57^. In all cases, however, this approach is critically limited as it treats the features extracted from different segments as *independent and identically distributed (i.i.d.)* observations from some underlying distributions. This independence is often implicit in the employed statistical and machine learning algorithms, but it ironically ignores mounting knowledge of the multiple-timescale dynamics of ictogenesis itself^58, 59^, as well as the various slow and ultra-slow dynamics of the circadian and multidien rhythms^5, 17, 21, 60, 61^, sleep-wake states^62, 63^, and seizure clustering^64, 65^ which all have well-documented modulatory effects on ictogenesis. As such, the standard approach misses a great opportunity for true “forecasting”, where the risk of a seizure occurring *s* minutes later is determined not based the brain’s current state, but based on what we predict its state to be *s* minutes into the future.

In this paper, we seek to improve the performance of existing machine learning-based seizure forecasting algorithms by augmenting them with two layers of predictive modeling: one at the level of iEEG features used as input for classification, and one at the level of seizure risk produced as output by the classification models. Our approach leverages the long-range temporal correlations in classification features to expand contextual information, and refines its estimates of seizure risk by filtering historical trends in its past risk estimates. Instead of proposing a new classifier, feature set, or model architecture, we introduce a general framework that operates independently of the specific choice of algorithm or features, adding additional layers to existing seizure prediction methods in a way that requires no retraining or fine-tuning of the baseline model. By integrating this framework with various baseline models, we demonstrate its capacity to consistently improve performance across a wide range of models, from support vector machines (SVMs) and random forests to convolutional neural networks (CNNs) and transformers.

## 2 Results

### Multi-level predictive modeling for improved seizure prediction

Significant prior work has pursued the design of seizure prediction algorithms from scalp and intracranial EEG^5–12^. Instead of seeking to design another such algorithm, in this work we take the orthogonal path of choosing a seizure prediction algorithm off the shelf and *augmenting it* with rich temporal statistical relationships that existing algorithms largely overlook. A typical seizure prediction/forecasting algorithm can be abstracted into a classifier *f* that maps a vector **x**[*t*] of features extracted from a moving window of (i)EEG up to time *t* (say, [*t* − *d, t*]) to the probability *q*_*B*_[*t*] that the given window is preictal (Figure 1):

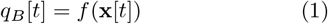

**Figure 1.**
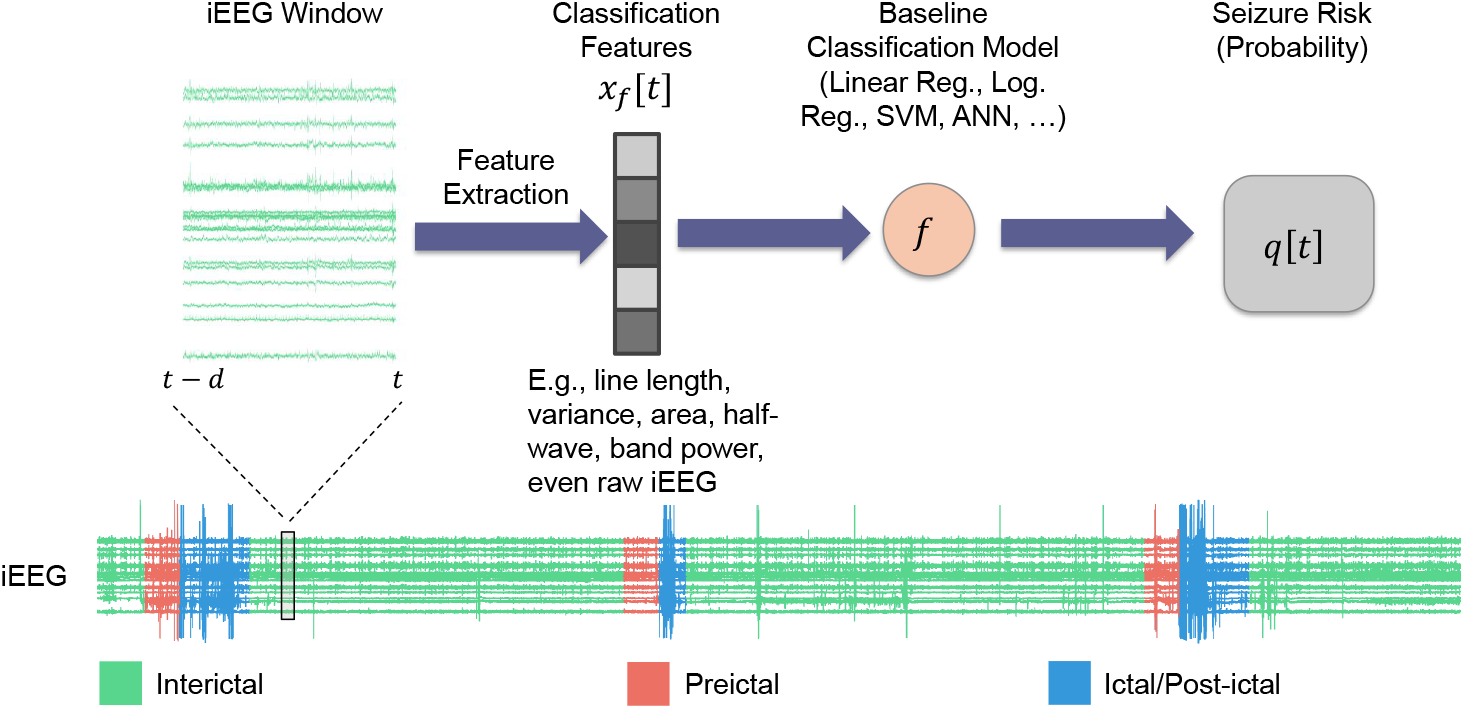
Standard seizure prediction pipeline. Existing seizure prediction/forecasting algorithms involve processing raw iEEG data through a series of steps to classify brain states as either preictal (seizure-impending) or interictal (baseline). This pipeline often starts with preprocessing to remove artifacts followed by extracting relevant features (**x**[*t*]), a classification model that uses these features to predict the preictal probability of the given window (*q*[*t*]), and an optional final thresholding to obtain a binary decision. While the same scheme may be used for any electrophysiological data, in this work we use pre-surgical iEEG from *n* = 5 patients obtained from the iEEG.org database (see Methods). In all of what follows, we compute features over windows of size *d* = 2 minutes, shifted each time by 1 step = 10 seconds.

In what follows we will refer to the function *f* as the *baseline model* and to time-varying probability *q*_*B*_[*t*] as the *seizure risk* at time *t* generated by this model. *q*_*B*_[*t*] may subsequently be thresholded to obtain a binary interictal/preictal decision if needed.

By nature, the above procedure is reactive rather than predictive. A positive (i.e., preictal) detection, even if made correctly, says something about a time window in the past (i.e., [*t* − *d, t*]) rather than a potentially impending seizure in the future. Therefore, we augment the above approach with two predictive, autoregressive dynamical models that are placed before and after the classifier, respectively (Figure 2). The former takes the general form

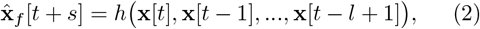

where *h* is a generally nonlinear dynamical model that takes as input the past *l* lags of classification features **x** from *t* − *l* + 1 to *t*, and predicts the future evolution of iEEG features *s* steps into the future. We tune the structure and hyper-parameters of *h* (particularly, *s* and *l*) and train Eq. (2) as a regression model to minimize the error between predicted 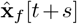 and true **x**[*t* + *s*] features over a patient-specific training set of data, separately for each patient and each feature (see Methods). Once learned, we feed the predictions of Eq. (2) at test time to the same classifier *f*, giving the predicted risk

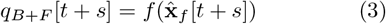

which is then used for pseudo-prospective seizure forecasting on a separate, within-patient test data (see Methods). The subscript _*B*+*F*_ reflects the addition of feature dynamics in Eq. (2) to the baseline classifier *f*, and distinguishes *q*_*B*+*F*_ from the *baseline risk* in Eq. (1).

**Figure 2.**
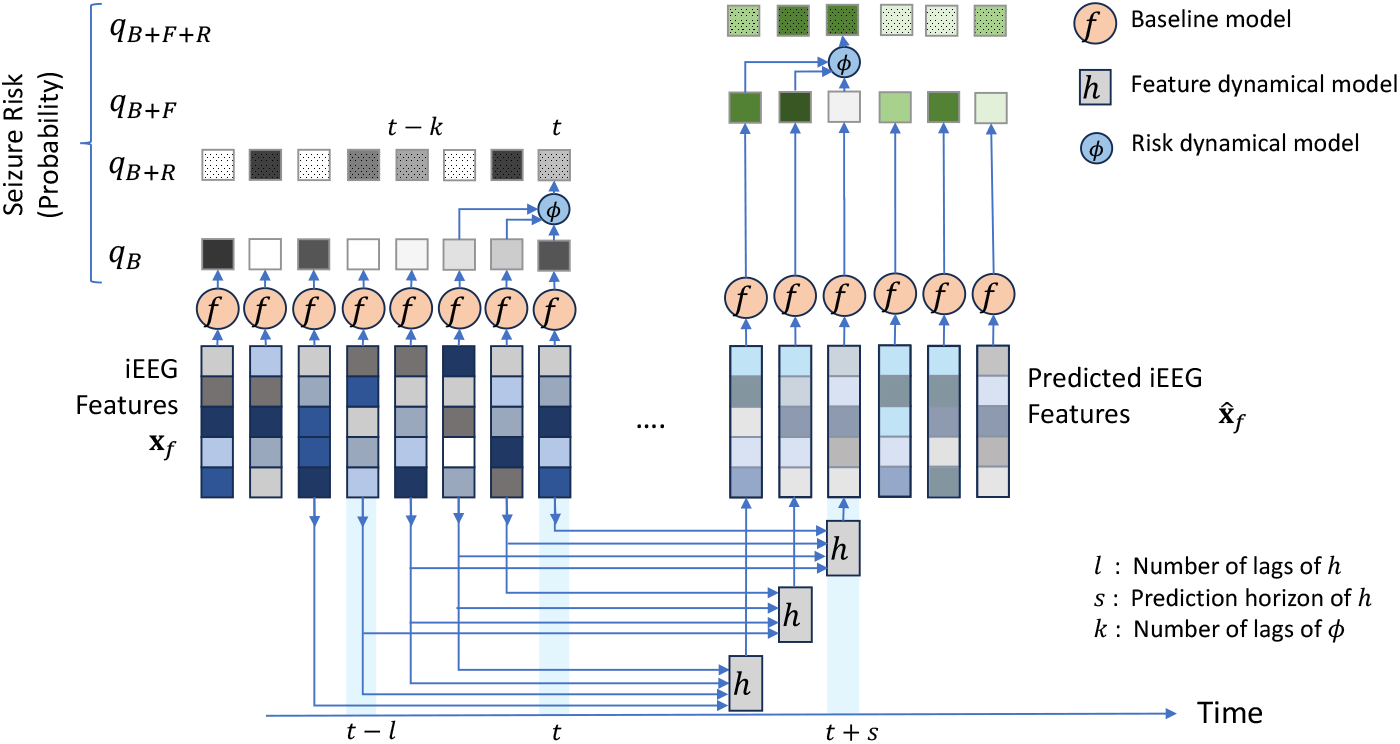
Proposed multi-level predictive modeling for improve seizure forecasting. Unlike the bulk of prior research, we start with an off-the-shelf seizure forecasting model *f* (cf. Figure 1) and augment it with one or both of two autoregressive predictive models: *h*, that operates on iEEG features (inputs to *f*), and *ϕ* that operates on seizure risk (outputs from *f*). All models and modeling choices, including the structure, parameters, and hyper-parameters of *h* and *ϕ* are tuned separately for each patient (see Methods). This results in 4 estimates of seizure risk *q*_*M*_, where *M* = *B* (baseline), *M* = *B* + *F* (baseline + feature dynamics), *M* = *B* + *R* (baseline + risk dynamics), or *M* = *B* + *F* + *R* (baseline + both dynamics). Each *q*_*M*_ is used for pseudo-prospective seizure forecasting on a held-out continuous test duration of each patient’s own iEEG.

A second autoregressive dynamical model is in turn built to capture temporal dynamics at the output of the classifier. This model generalizes the commonsense approach of looking at a window *q*_*B*_[*t* − *k* + 1], …, *q*_*B*_[*t*] of seizure risks to make a interictal/preictal decision, and similar to Eq. (2) takes the general form (Figure 2)

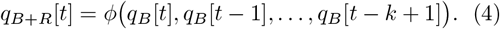

The function *ϕ* and the lag hyper-parameter *k* are trained separately for each patient (see Methods). Given that each *q*_*B*_[*t* − *τ*] term is dependent on the underlying feature **x**[*t* − *τ*], this approach can be also considered a way to indirectly increase the iEEG context available for seizure forecasting and to refine the seizure risk *q*_*B*_[*t*] at time *t* based on its historical trends. Finally, we allow for the combination of feature and risk dynamics, giving rise to the combined estimate

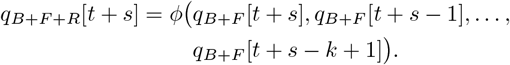

In general, the choice of dynamical model (feature, risk, or combined) that leads to best pseudoprospective accuracy varies from patient to patient and needs to be optimized per patient.

### iEEG features follow predictable dynamics up to 30 minutes into the future

We first investigated the temporal predictability of various iEEG-based features that are commonly used for seizure forecasting. This is important because at the core of the predictive scheme described above (Figure 2) is the autoregressive model in Eq. (2) that predicts the evolution of iEEG features into the future.^†^ The viability of our scheme is therefore contingent on the presence of predictable dynamics in iEEG features **x**[*t*].

Using an autoregressive (AR) form for feature dynamics as in Eq. (2), we examined how accurately each iEEG feature can be predicted into the future, with a focus on assessing whether features remained predictable over extended periods. In general the AR function *h* can (and often should, cf. Figure 5) be nonlinear, but for this analysis we limited *h* to be linear for simplicity and computational feasibility. We then parametrically varied the horizon length *s* for every patient and every feature, and measured crossvalidated (out-of-sample) regression accuracy (*R*^2^) as a function of *s*.

The results, as shown in Figure 3a, display the distribution of regression *R*^2^ values as a function prediction horizon *s*, combined across a wide range of seizure-relevant iEEG features separately for each patient (see Methods). As expected, the predictive accuracy generally decreases with prediction horizon *s*. Interestingly, however, most features maintained above-chance predictability (*R*^2^ *>* 0) even over long horizons up to 10 minutes. Beyond the 10-minute mark more than half of the features often exhibit a negative *R*^2^ (lower than chance predictability). Yet, in each patient there are some features that remain predictable even up to 30 minutes. These results support the hypothesis that iEEG features hold intrinsic temporal structures that persist over long horizons, providing a basis for long-term seizure forecasting.

**Figure 3.**
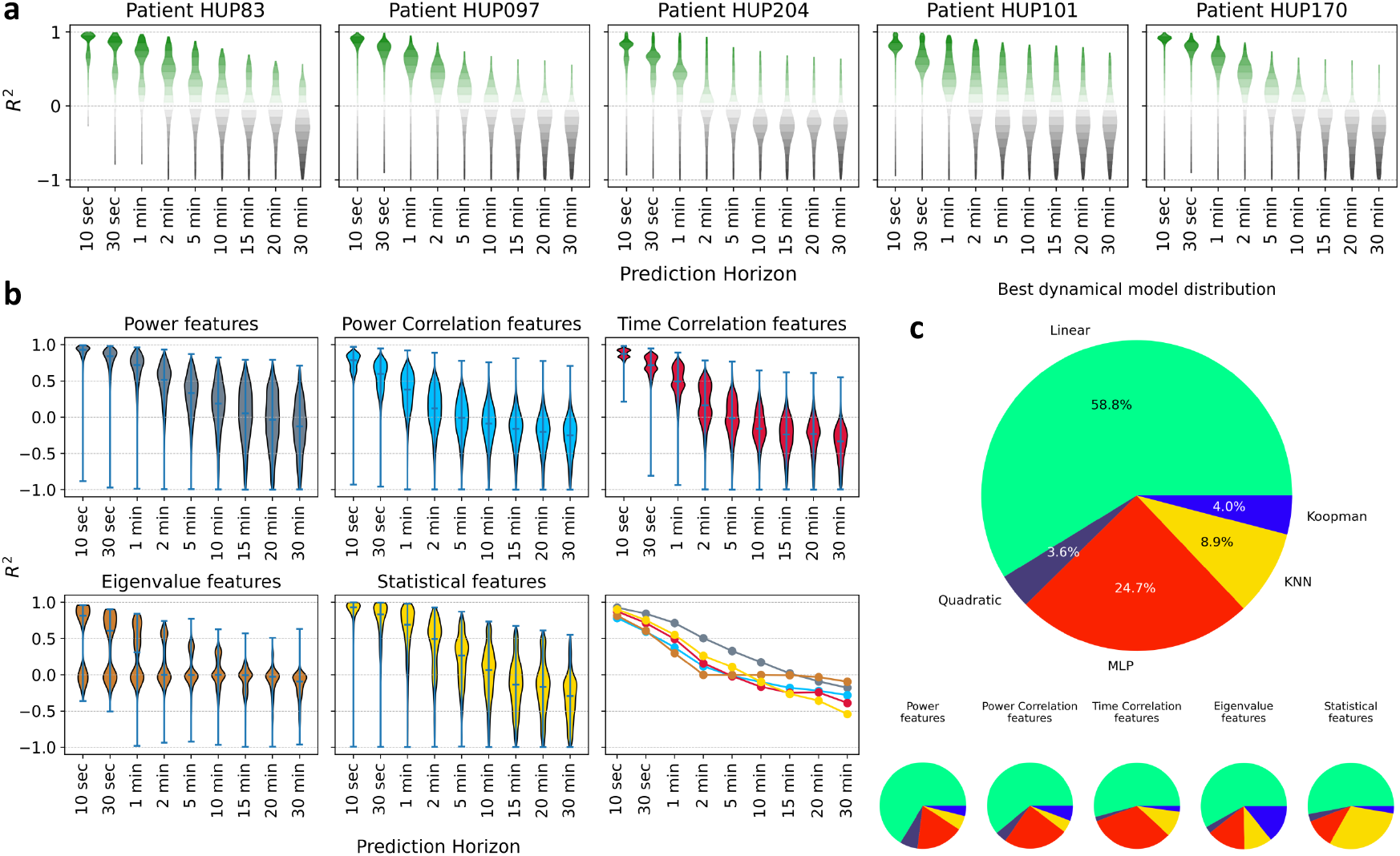
Long-horizon predictability of seizure-relevant iEEG features using autoregressive (AR) dynamical models. **(a)** Each panel, corresponding to one patient indicated at the top, shows the distribution of cross-validated *R*^2^ of a linear *s*-step-ahead AR model as in Eq. (2) as a function of prediction horizon *s*. The vertical range in all panels is limited to [−1, 1] for better visualization. Each violin plot is combined across all features, which are themselves a union of various sets of features used by different seizure classification models in the literature (see Table 2 for a categorical breakdown and Methods for details). Green (resp., gray) indicates positive (negative) *R*^2^ and better (worse) than chance performance. Most features remain predictable above chance up to 2-10 minutes depending on the patient, and across all patients some features remain predictable above chance up to 30 minutes. **(b)** Predictability of iEEG features by category. Data is the same as panel (a) but now combined across patients and separated by feature category (average number of features across patients: power features ≃2800, power correlation features ≃8000, time correlation features ≃8000, statistical features ≃1600, eigenvalue features ≃250). For a complete list of features in each category, see Methods. The bottom right sub-panel shows the medians of all *R*^2^ distributions overlaid and color-coded by feature category. Power features clearly have the longest predictability compared to other categories. **(c)** Distribution of AR models that explain most variance (have maximum *R*^2^) across all features and patients. For each patient-feature, we fit five different AR models (1 linear, 4 nonlinear) and selected the one with the highest cross-validated *R*^2^. MLP: multi-layer perceptron, KNN: k-nearest neighbor. See Methods for details on each model.

Importantly, the long-horizon predictability of iEEG features varies significantly across feature categories (time-domain, frequency-domain, etc.). As shown in Figure 3b, power features have the farthest predictability, while the predictability of eigenvalue features (eigenvalues of channel cross-correlation matrices) drops to chance level most quickly (see Methods for details). Yet, not all frequency-domain features are necessarily better predictable than time-domain features; power correlation features, e.g., decay to chance-level more quickly than time correlation features. Though beyond the scope of this work, these differences in predictability can be very valuable for improved feature engineering and classifier design for seizure forecasting.

Further, using potentially nonlinear AR models can further improve the long-horizon predictability of iEEG features, but only marginally. For each patient and iEEG feature, we compared five different functional forms for Eq. (2)–the linear and 4 nonlinear forms. We then found the most accurate model for each patient-feature and computed the win percentage of model across all patient-features. As shown in Figure 3c, in about three quarter of cases, the linear model achieves the highest *R*^2^, followed by the multi-layer perceptron artificial neural network that approximately covers the remaining quarter of patient-features. Notably, this result shows that the dynamic linearity we had found earlier at the level of raw iEEG potentials^66^ *approximately* holds also for a wide range of iEEG features. Though weak, however, the nonlinearity of iEEG features is not negligible and will in fact prove to be critical for dynamics-augmented seizure forecasting (cf. Figure 5).

### iEEG features with higher predictability tend to also have higher importance in seizure classification

We next examined the importance of each iEEG feature for seizure classification across several state-of-the-art seizure forecasting models, and compared the importance of each feature with its long-horizon predictability (Figure 4). While the exact definition of feature importance varies be-tween classification models (see Methods), it generally quantifies the sensitivity of the classifier’s output to variations in each input (feature) and is a metric of how relevant each feature has been for seizure classification based on training data. Note that, in general, the importance and predictability of features are independent quantities coming from two orthogonal approaches to the iEEG time series (cross-sectional seizure-focused classification vs. temporal seizure-independent regression). Therefore, the relationship between feature importance and feature predictability is critical for the success of integrating feature dynamics (Eq. (2)) into seizure forecasting. In particular, it is the *conjunction* of feature importance and predictability that will make dynamics-augmenting potentially beneficial, whereas either alone is unlikely to be sufficient.

**Figure 4.**
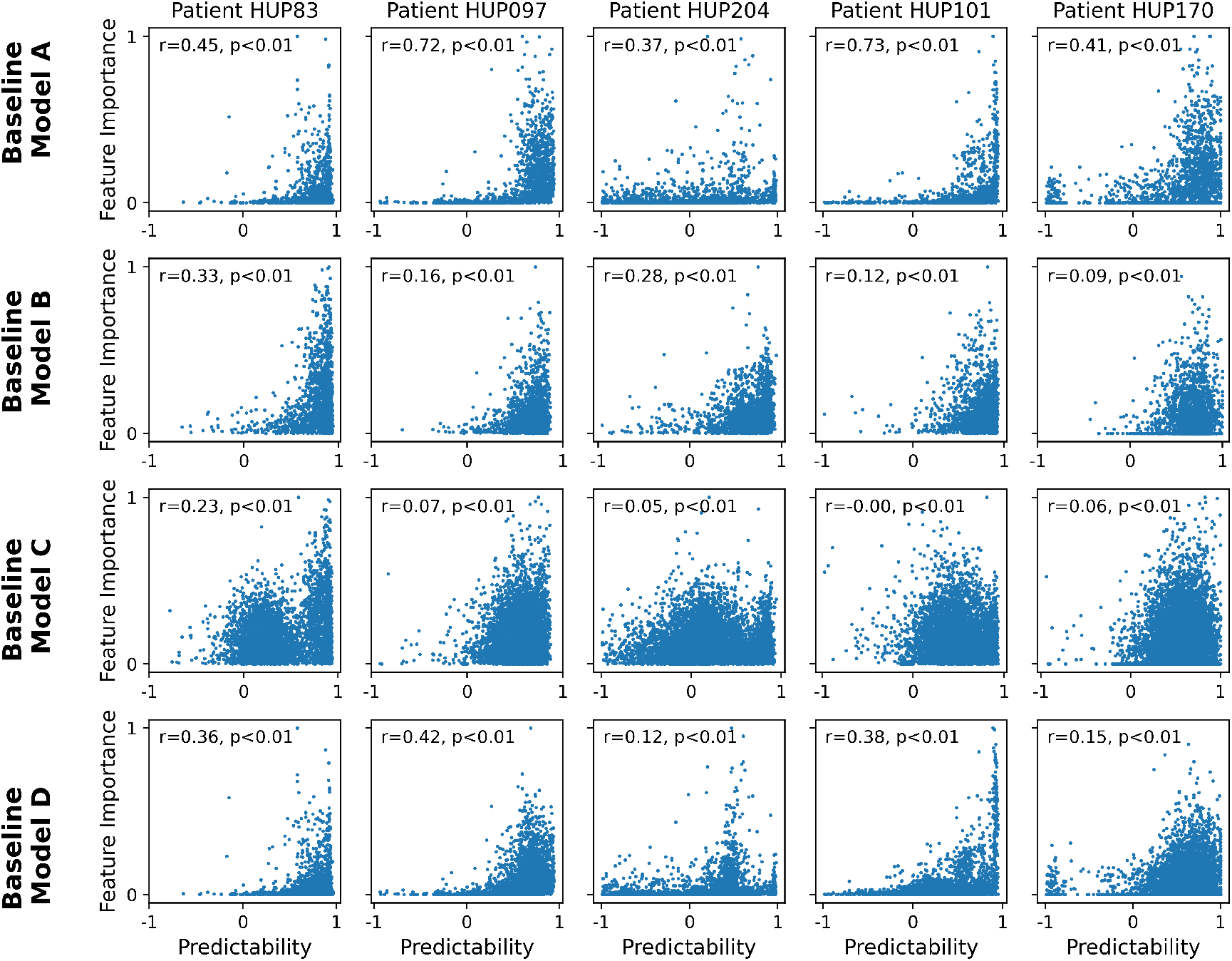
Presence of significant positive correlation between feature predictability and feature importance across various state-of-the-art seizure classification models. Each panel shows the relationship between feature predictability and feature importance for one patient and one classification model. Each dot represents one iEEG feature, and predictability is computed as the cross-validated *R*^2^ of a linear AR model with a prediction horizon (*s*) of 2 minutes. The horizontal range of each panel is limited to [− 1, 1] for better visualization. Feature importance is computed based on the respective baseline model, see Methods for details. The Spearman correlation coefficient and associated p-value are shown in each panel.

As seen from Figure 4, in nearly all baseline models and all patients, there exists a strong positive correlation between feature importance and feature predictability. As with most aspects of seizure forecasting, there exists strong heterogeneity among patients as well as baseline models. In general, the positive correlation between feature importance and predictability is strongest for baseline model B and weakest for the baseline model C. Further, except for model C, this relationship is often generally monotonic and approximately exponential, with a rapid and strong increase in feature importance as predictability approaches 1. These results thus strongly support the potential benefit of dynamics-augmentation for seizure forecasting (Figure 2), as we directly examine next.

### Incorporating feature dynamics improves accuracy of pseudo-prospective seizure forecasting

Building on the established predictability of seizure-relevant iEEG features over extended time intervals, we used the estimated seizure risk *q*_*B*+*F*_ in Eq. (3) for continuous pseudo-prospective seizure forecasting using iEEG data from five patients from the Hospital of the University of Pennsylvania (HUP) on the iEEG.org portal. To demonstrate that our approach is robust and can adapt to various baseline models, we implemented six different state-of-the-art seizure forecasting models as the baseline model (Figure 1 and Eq. (1)) for each patient (see Methods). These models reflect a wide range of effective strategies in seizure forecasting, including models recognized for their strong performance in seizure research (Baseline Models A and B), models that have excelled in Kaggle-hosted crowed-sourced competitions (Models C and D), and approaches using state-of-theart machine learning (Models E and F). In each case, we measure seizure forecasting accuracy based on the area under the pseudo-prospective sensitivity-time in warning curve (PP-AUC, Figure 8, see Methods for details).

The results, summarized in Figure 5, demonstrate a strong improvement in seizure forecasting accuracy across nearly all patients and baseline models. For each patient-baseline combination, we evaluated the same five candidates for feature dynamical modeling (*h* in Eq. (2)) as in Figure 3c and selected the one with the highest PP-AUC. Figure 5a shows a breakdown of average PP-AUC for baseline and best-performing dynamics-augmented alternative across all patients and baseline models. The success rate of each dynamical model and the distribution of relative improvement in PP-AUC are summarized in panels b and c, respectively. Except for one case where the baseline model alone is slightly better than all dynamical alternatives, in all other patient-baseline combinations augmenting the baseline model with AR feature dynamics results in a higher PP-AUC. Across all patient-baseline combinations, we obtain a mean increase in PP-AUC of 28%, signifying the substantial but untapped potential of simple AR modeling for improved seizure forecasting.

**Figure 5.**
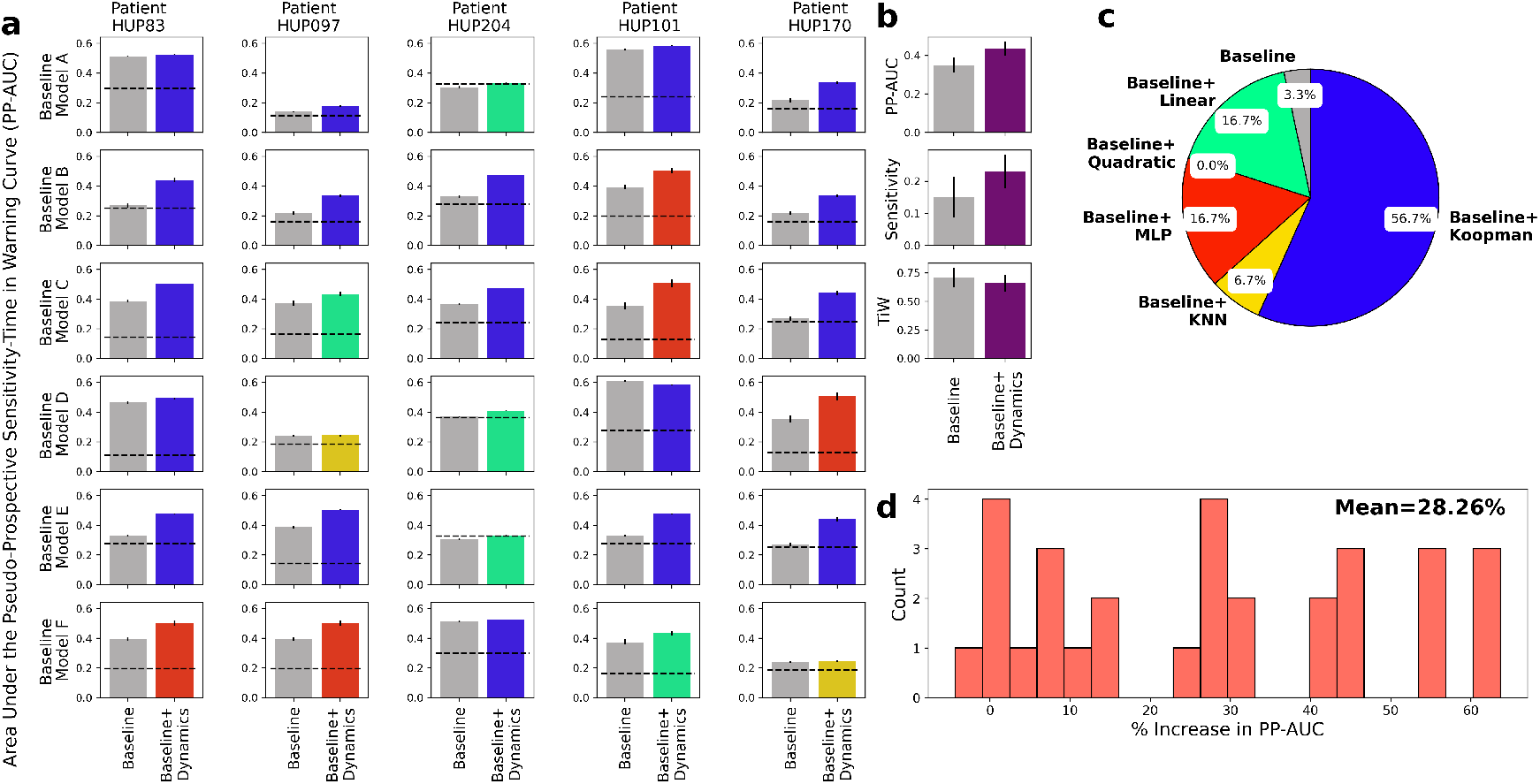
Pseudo-prospective seizure forecasting accuracy improves after incorporating predictive dynamical modeling of iEEG features. **(a)** Area under the pseudo-prospective sensitivity-time in warning curve (PP-AUC, cf. Figure 8 and Methods for details) for the baseline model (left, gray) and best dynamics-augmented alternative (right, colored) across five patients and six baseline models. The color of the right bar in each panel illustrates the best-performing model *h* and matches the color code in (b). Each bar shows mean *±* 2 s.e.m. over randomized train-test splits. **(b)** Summarizing the results in panel (a) across patients and baseline models. The use of dynamics results in higher PP-AUC (top), higher sensitivity at 30% time-in-warning (TiW) (middle), and lower TiW at 60% sensitivity (bottom). **(c)** Distribution of AR models *h* that achieved the highest PP-AUC across all patients and baseline models. Note the major differences with the similar distribution in Figure 3. **(d)** Distribution (histogram) of the relative improvement in PP-AUC across all patient-baseline combinations in (a). Relative improvement is measured as (PP-AUC_B+D_ -PP-AUC_B_) / PP-AUC_B_.

Besides improving the accuracy of seizure forecasting, predictive dynamical models can provide insights into the nature of seizure dynamics. In particular, the optimal length of “past” and “future” that maximizes seizure forecasting accuracy are important markers of the timescale at which seizures arise in each patient (Figure 6). Note that these values are based on optimizing seizure forecasting accuracy (PP-AUC) and are thus different from *R*^2^ values shown in Figure 3 that reflect pure feature autoregressive accuracy. Notably, we can see from Figure 6 that both the optimal number of autoregressive lags (past) and prediction horizons (future) are internally stable for each patient, externally variable across patients, and overall on the order of ∼ 10 minutes for all patients. These values, while consistent with the earliest reports of the preictal period^4^, provide detailed measures of temporal structures in iEEG that are sufficiently robust and reproducible in each patient that can be used for automated seizure forecasting.

**Figure 6.**
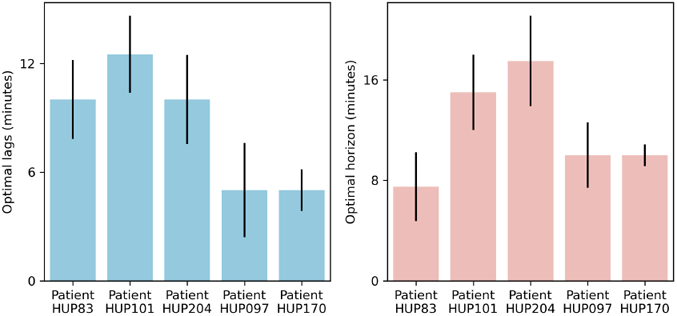
Optimal temporal timescale of iEEG feature dynamics for seizure forecasting. The left and right panels show the optimal number of autoregressive lags (history length) and the optimal prediction horizon (step size), respectively. Both values are obtained using the linear *h* model based on maximizing the PP-AUC score and reported separately per patient as median (across all features for that patient) ± 1 s.e.m. Note that both the optimal number of lags and the optimal prediction horizon varies considerably across patients, reflecting the vast heterogeneities that exist among epilepsy patients and the need for patient-specific modeling.

### Modeling risk dynamics further enhances seizure predictability

Next we examined the effects of adding a second layer of predictive dynamics at the level of seizure risk, Eq. (4), on pseudoprospective seizure forecasting. Similar to above, we for each patient-baseline combination we used a training portion of iEEG data to fit the baseline model, five feature-predictive models *h*, and one risk-predictive model *ϕ*, resulting in 12 possible combinations (baseline alone or baseline with either of the 5 forms of *h*, each with or without *ϕ*). We then evaluated the PP-AUC of each of these 12 possibilities in patient-specific test data, repeated over all train-test splits, and found the model with the highest mean PP-AUC for each patient-baseline combination.

The results, shown in Figure 7, demonstrate the additional benefit of including risk dynamics over not only the baseline but also the combination of baseline and feature dynamics. Similar to Figure 5, inclusion of dynamic predictive models improves PP-AUC in all except one patient-baseline combination (Figure 7a). However, the addition of risk dynamics has further improved PP-AUC in half of the cases (Figure 7c), bringing the total mean improvement in PP-AUC over baseline to 51% (Figure 7d).

**Figure 7.**
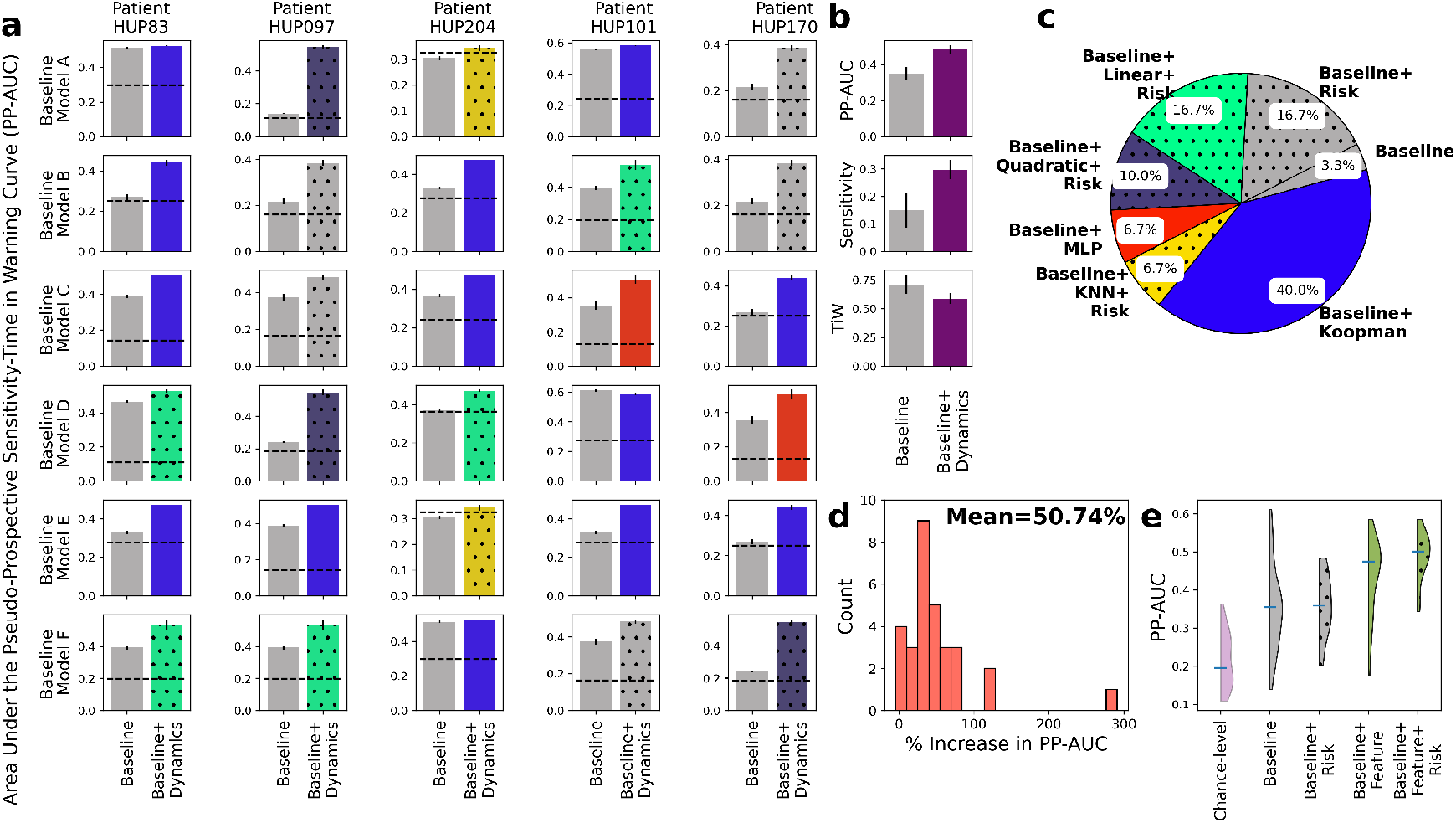
Further improvements in seizure forecasting accuracy with incorporation of feature *and risk* dynamics. **(a)** Side-by-side breakdown of improvements in PP-AUC resulting from incorporation of both feature (Eq. (2)) and risk (Eq. (4)) dynamics. Details are similar to Figure 5a. In each sub-panel, the color of the bar on the right still shows the form of feature dynamics *h* that resulted in highest PP-AUC, and the dotted hatches show the presence of risk dynamics in the best-performing model. **(b)** Summary comparisons of PP-AUC, sensitivity at 30% TiW, and TiW at 60% sensitivity between baseline and dynamics-augmented forecasters across all patients and baseline models. Details parallel those in Figure 5b. **(c)** Distribution of models that achieved the highest PP-AUC across all patients and baseline models. Details are similar to Figure 5c with the addition of dotted hatches showing the presence of risk dynamics in the best-performing model as in (a). **(d)** Distribution of relative improvement in PP-AUC across all patient-baseline combinations similar to Figure 5d. **(e)** Distributions of raw PP-AUCs for all four possible combinations of baseline models with feature and/or risk dynamics. Each distribution combines across all 5 patients and 6 baseline models.

**Figure 8.**
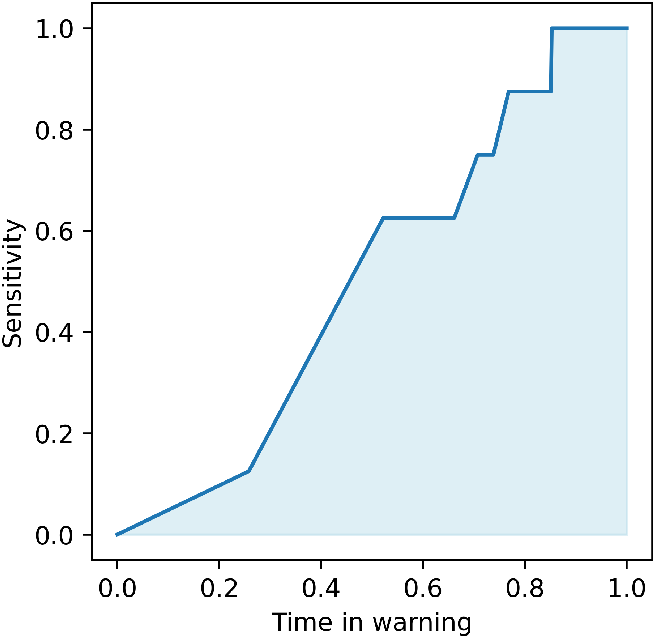
Illustrative example of performance metric used for comparison between methods, namely, the pseudo-prospective area under the curve (PP-AUC). This metric closely resembles the standard area under the receiving operator characteristic curve, with the main difference being the substitution of false positive rate with time in warning.

Figure 7e shows the overall distribution of PP-AUC across all patient-baseline combinations using *q*_*B*_, *q*_*B*+*F*_, *q*_*B*+*R*_, and *q*_*B*+*F* +*R*_. The addition of risk dynamics, both in the absence and presence of feature dynamics, plays a more regulatory role, reducing outliers and increasing the robustness and reliability of seizure forecasting via a dynamic filtering of the risk dynamics. This effect can also be seen from the detailed breakdown of seizure forecasting with and without the risk dynamics alone (Supplementary Figure 1) and results in only a marginal gain in the absence of feature prediction. It is the latter that is the main driver of improved PP-AUC, and the combination of the two dynamic layers that results in best seizure forecasting overall.

## 3 Discussions

Seizure prediction and forecasting research over the past two decades has largely reduced these problems to statistical classification and/or regression, and relied primarily on advancements in machine learning and data acquisition to improve outcomes. In this study we provided pioneering evidence for the utility of an orthogonal dimension of improvement, i.e., explicitly modeling temporal dynamics over classification features and seizure risk. Using publically available pre-surgical iEEG data from five patients at the Hospital of the University of Pennsylvania (HUP) and six state-of-the-art seizure forecasting models as baseline, we showed that adding simple autoregressive (AR) predictive models at the level of iEEG features can increase pseudo-prospective forecasting accuracy by an average of 28%. This improvement increased to 51% when we added a second AR model at the level of seizure risk, while the latter alone had marginal benefit. Additional analyses revealed that these benefits primarily stem from the convergence of two factors: long-term predictability of many iEEG features up to tens of minutes, and strong positive correlation between the predictability of iEEG features and their importance for seizure classification. In addition to revealing a new dimension of temporal dynamics in the context of ictogenesis, these results open the door to various future studies into the benefits of predictive modeling in specific applications of seizure prediction and forecasting.

In general, the benefits of predictive modeling in a context such as seizure forecasting depends on a trade off between accuracy and latency. On the one hand, predictive modeling (Eq. (2)) uses temporal statistical correlations to predict iEEG features several minutes before they occur, and this creates an advantage in terms of latency. However, on the other hand, any predictive model will inevitably introduce errors due to the stochastic nature of the underlying time series, suboptimal modeling, etc. This creates a trade off (which becomes stronger as the prediction horizon *s* increase) between using “future” features which are only approximate, or using present features which are perfectly accurate. As such, it is neither trivial nor guaranteed that adding predictive modeling should improve seizure forecasting, and which effect dominates is in general patient-, baseline classifier-, and horizon-dependent. Given this context, our findings are significant as they show that for almost all patient-baseline combinations and at least some prediction horizons, it is latency that dominates accuracy.

Our study further addresses an important gap in seizure forecasting research, namely, moving beyond curated and pre-segmented preictal-interictal data and benchmarking algorithms based on pseudoprospective analysis (PPA) on continuous data. As clearly seen from Figure 7e, the median pseudoprospective AUC across all the models and patients we tested is about 0.35 at baseline (and about 0.5 even after addition of predictive modeling). This is alarmingly lower than commonly reported AUCs on the basis of curated data (see, e.g., Ref.^67^). The latter values are likely to be overly optimistic and not the most accurate reflection of how each algorithm would actually perform in a clinical setting. One major reason behind this gap is the *additional difficulty of preventing false positives* when an algorithm is monitoring continuous iEEG with long interictal durations which can contain numerous “seizure-looking” motifs.

This is a serious challenge, however, and in need of more focused attention in future research. Whether a seizure forecasting algorithm is used to deliver advisory warnings to the patient or closed-loop neurostimulation to the brain, the additional challenges of PPA are crucial and inevitable in either setting.

In the context of PPA, also notable are the differences between acutely- and chronically-recorded iEEG. In this study we used pre-surgical iEEG data recorded acutely at the HUP epilepsy monitoring unit (EMU). By nature, however, EMU conditions are highly non-stationary and recorded seizures are likely more heterogeneous in their mechanisms and dynamics than each patient’s normal seizures outside of the hospital. The number of recorded seizures is also very limited in acute data, with only a few for most patients and hardly over a dozen in any patient. Both factors (abnormally high heterogeneity and very few recorded seizures) are major bottlenecks in using acute data, particularly for data-driven algorithms like those investigated here. We therefore expect seizure forecasting algorithms to reach (potentially significantly) higher PP-AUCs if used with chronically-recorded data. On the other hand, however, chronically recorded iEEG has significantly lower spatiotemporal coverage than acute iEEG, and future work is needed to rigorously test and benchmark the algorithms studied in this work on chronically recorded data. Also a promising avenue for future research is testing the benefits of predictive modeling on seizure forecasting using scalp EEG and even other physiological time series (heart rate, seizure occurrence, etc.).

A long-standing question in epilepsy research is what exactly constitutes the “preictal” period^68^. In this work we initially used the common one-hour preseizure window (precisely, 65-5min before seizure onset) for labeling data for the purpose of training each classifier. However, our predictive modeling results yielded optimal prediction horizons (i.e., value *s*^*^ that resulted in maximum PP-AUC) that varied widely across patients, ranging approximately between 7 to 20 minutes (Figure 6). These values are interestingly consistent with the earliest reports of the preictal period^4^, and are promising from a clinical standpoint as even a five-minute warning could give patients enough time to reach a safe space or administer medication to reduce the risk of seizure occurrence. At the same time, these results highlight the importance of accurate patient-specific characterization of the preictal period, and suggest *s*^*^ as a potential data-driven biomarker that can be used for this purpose.

An interesting finding of our study was a major disagreement between the choice of predictive models of iEEG features that directly maximized regression *R*^2^ (Figure 3c) and the choice that maximized downstream pseudo-prospective forecasting accuracy (Figure 5c). As far as the former is concerned, a linear AR model best described iEEG feature dynamics in most cases. However, complex, nonlinear models (particularly those based on the Koopman operator theory) led to greater accuracy when used in the context of seizure forecasting. This is interesting from at least two perspectives. First is the importance of end-to-end (rather than piece-by-piece) optimization when multiple analyses/models are combined within a larger pipeline. Second is the additional complexity of seizure forecasting compared to pure iEEG feature prediction. This additional complexity may be the reason why a linear model can be optimal for pure feature prediction, but a more expressive nonlinear model may give predictions that are more useful for seizure forecasting. Future work is needed to dive more deeply into and mechanistically explain the na-ture of the dynamics of iEEG features and seizure risk.

This study has a number of limitations. The exploratory nature of our work required us to implement and test a combinatorially large number of structural and parametric choices for each patient, each of which required running computationally expensive algorithms on continuous iEEG from tens to hundreds of channels over days to weeks of recording. Therefore, in this work we only used data from 5 patients that had a (relatively) large numbers of lead seizures (Table 1). In this selection we did not further restrict the patient’s type of epilepsy in order to maximize the number of lead seizures per patient, a choice that also showcases the promise of our approach across diverse epilepsies. Future work will aim to extend our findings to larger cohorts through more focused, hypothesis-driven investigations. Another limitation of this work is that our dynamical models, similar to the baseline classifiers they augment, are data-driven and do not provide a mechanistic understanding of the relationship between each individual’s pathophysiology and their predictive dynamics. This, in particular, makes it harder to generalize models across individuals and limits the success of seizure forecasting to the quality and quantity of available data. Another constraint is that seizure annotations that we have used (i.e., “the ground truth”) were have been performed manually by a board-certified epileptologist, limiting our analyses to seizures that have been documented in clinical records. Human errors and inter-rater variability have been documented and studied extensively, but manual annotations still remain the gold standard due largely to the complexities of automated seizure detection^69, 70^. Finally, our current implementation does not impose limitations on CPU or memory usage. This flexibility is advantageous for exploratory work but will need to be addressed for future translations to implantable devices.

**Table 1:** Data characteristics of the patients used in this study. “Other cortex” refers to seizure onset zones outside the mesial temporal or temporal neocortical structures.

| Patient ID | Seizure Onset Zone | # Channels | Data duration (days) | # Lead Seizures |
| --- | --- | --- | --- | --- |
| HUP083 | Other Cortex | 92 | 13.7 | 11 |
| HUP097 | Multifocal | 102 | 13.1 | 12 |
| HUP101 | Multifocal | 102 | 12.3 | 10 |
| HUP170 | Other Cortex | 228 | 6.7 | 13 |
| HUP204 | Mesial Temporal Lobe | 117 | 6.1 | 10 |

In sum, our findings underscore the importance of slow dynamics in seizure forecasting, provide an innovative approach for using slow dynamics at the levels of iEEG features and seizure risk, and illustrate a largely-untapped potential for improved seizure forecasting. These insights pave the way for future work focused on enhancing model robustness and generalizability, meta-learning across large cohorts of patients, and prospective clinical testing in both acute and chronic contexts.

## 4 Methods

### Data

All analyses and experiments presented in this paper were conducted using publicly available iEEG data from the International Epilepsy Electrophysiology Portal (iEEG.org). The data for all patients consisted of stereoelectroencephalography (sEEG) recordings. We present results on *n* = 5 patients, each with varying seizure characteristics, electrode channels, and seizure durations, as summarized in Table 1. patient selection was based on the availability of a sufficient number of lead seizures (≥ 10), which is critical for enabling a reliable train-test split in the seizure prediction framework. While additional patients were available in the database, only a small subset satisfied this criterion, and we further restricted our analyses to five patients in order to manage the extensive computational experiments required for this study. We down-sampled all recordings to 200 Hz for uniformity and lowering the computation complexity of feature extraction. For each patient, seizure onset and offset times were manually annotated by a trained epileptologist and accompanied the raw data on iEEG.org.

Using the inter-seizure intervals we annotated a seizure as a lead seizure if it occurred at least 8 hours away from the previous seizure offset. Interictal periods were defined as durations occurring at least 3 hours away from any seizure event in both temporal directions, whereas the interval [65, 5] minutes before each lead seizure was labeled as preictal. For each patient, we constructed the training set using the preictal periods from the first three lead seizures, together with three neighboring interictal periods of 1 hour each. To further ensure that the baseline classifier had sufficient training data, we also randomly selected one or two additional lead seizures (with matched interictal segments) for inclusion in the training set. Limiting this number to at most two strikes a balance between providing enough data for classifier training and retaining the majority of seizures for unbiased pseudo-prospective evaluation. The corresponding segments were excluded from testing, and all remaining iEEG data after the third lead seizure were reserved for evaluation.

### Baseline classification models

As noted earlier, throughout this work we started from a given (off-the-shelf) “baseline” seizure forecasting model (Eq. (1)) and studied the benefits of augmenting it with predictive models at the level of its input (iEEG features, **x**) and/or output (seizure risk, *q*). Therefore, our approach is in general agnostic to the choice of baseline model *f* and its iEEG features **x**_*f*_, and we have evaluated it using a diverse set of baseline models and features. Specifically, we used six different baseline models, as summarized in Table 2. We selected these models based on the following criteria: those employed in influential papers on seizure forecasting (Models A and B), those shown to be effective in crowd-sourced Kaggle competitions (Models C and D), and those representative of modern machine learning approaches (Models E and F). For the latter four, publicly available code facilitated implementation, whereas for the first two, we replicated their classifiers using our custom code based on our best understanding of the corresponding papers and their available supplementary material.

**Table 2:** List of baseline seizure forecasting models used in this work.

| Baseline Model | Classifier ($f$) | Features ($\mathbf{x}$) | Reference |
| --- | --- | --- | --- |
| A | k-Nearest Neighbor | Power features, Statistical features (line-length, signal average) | Based on Ref.<sup>71</sup> |
| B | Logistic Regression | Power features | Based on Ref.<sup>72</sup> |
| C | Support Vector Machine | Power features, Power correlation, Eigenvalues | ‘Birchwood’ model in Ref.<sup>73</sup> |
| D | Random Forest | Power features, Time correlation, Eigenvalues, Statistical features (channel kurtosis, standard deviation, line-length) | Team B in Ref.<sup>15</sup> |
| E | Convolutional Neural Network | Power features | Based on Ref.<sup>47</sup> |
| F | Transformer | Power feature | Model in Ref.<sup>57</sup> |

For each baseline model we computed its respective features as originally proposed on a 2-minute moving window. The window was shifted each time by 50ms and 10s on training and test data, respectively. After feature extraction, we train each baseline model as a supervised binary classifier and extract its resective seizure risk 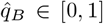 from before the final thresholding step. The resulting, continuous seizure risk trajectory 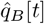 was computed on both the training and test samples. The former was used to train the risk-predictive models in Eq. (4), while the latter were used to compute each baseline model’s PP-AUC (see below).

### Feature categories

The features used for model training were grouped into five categories to better illustrate their differences in dynamics (Figure 3). *Power features* capture the spectral content of the iEEG by computing band-limited power for each channel, either in canonical frequency bands (delta, theta, alpha, beta, gamma) or in equally spaced frequency bins used in Baseline Models C. In total, for each patient we had 23*C* power features, where *C* denotes the number of iEEG channels (see Table 1). *Power correlation features* measure the interdependence of spectral activity across channels by computing correlation matrices between band power time series, e.g., correlating the power in band 1 of channel 12 with the power in band 3 of channel 4. The total number of power correlation features equals*C ×* (*C*− 1)*/*2. for each patient. *Time correlation features* similarly compute pairwise correlations, but directly on the raw iEEG time series rather than on band powers, quantifying linear relationships between channels in the time domain. Similar to power correlation features, for each patient the total number of time correlation features is given by *C ×* (*C*− 1)*/*2. *Eigenvalue features* summarize the global structure of these correlation matrices by using their eigenvalues (2*C* features in total), which capture overall network synchrony and connectivity patterns. Finally, *Statistical features* are various simple time-domain signal statistics from each channel, which may vary slightly across baseline models. Typical examples include line length, mean, standard deviation, and the standard deviation of first differences. These features capture basic amplitude and variability characteristics of the signals. In total we had 13*C* statistical features for each patient.

### Computation of feature importance for baseline models

To relate feature predictability (from the feature dynamical model) to their relevance for classification, we computed feature importance scores for the baseline classifiers using model-appropriate approaches. For *k*-nearest neighbors (Model A), where importance is not explicitly defined, we used a permutation approach: feature values were shuffled across samples, and the resulting drop in accuracy was recorded. For logistic regression and linear SVM (Models B and C), importance was taken as the absolute value of the learned coefficients or weight vector, reflecting the strength of association between features and the decision boundary. For the random forest (Model D), importance was quantified by the mean decrease in Gini impurity, as implemented by the `scikit-learn` package in python. For the deep learning models (Models E and F), we did not compute feature importance, since attribution methods (e.g., saliency maps) are computationally intensive and beyond the scope of this study.

### Predictive modeling of iEEG feature dynamics

For each patient we learned the feature-predictive model *h* in Eq. (2) as a nonlinear regression problem. We considered five different forms for *h*, as detailed in Table 3, and tuned the parameters ***θ*** of each model by minimizing its sum of squared error

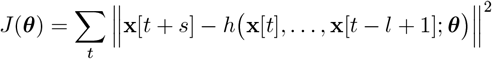

over the training dataset. In all cases, the function *h* was learned element by element, such that the future values of each iEEG feature were predicted us-ing its own past history. The hyper-parameters of each model were fixed as listed in Table 3, except for the main hyper-parameters *l* and *s*, which were optimized by iterating over *l* ∈ {5, 10, 15 } min and *s*∈ { 1*/*6, 1*/*2, 1, 2, 5, 10, 15, 20, 30} min separately for each patient.

**Table 3:** List of autoregressive dynamical models used for *s*-step ahead prediction of iEEG features in Eq. (2). KNN: k-nearest neighbor; MLP: multi-layer perceptron.

| Model | Equation |
| --- | --- |
| Linear | $\hat{x}_i[t + s] = \sum_{n=0}^{l-1} \alpha_n x_i[t - n]$ |
| Quadratic | $\hat{x}_i[t + s] = \sum_{m,n=0}^{l-1} \beta_{m,n} x_i[t - m] x_i[t - n]$ |
| KNN | $\hat{x}_i[t + s] = \frac{1}{K} \sum_{n=0}^{K-1} x_i^{\text{train}}[t_n + s]$ where $t_0, \dots, t_{K-1}$ are time indices of the $K$-nearest neighbors among training data to $(x_i[t - l + 1], \dots, x_i[t])$. We used $K = 5$ for all patients. |
| MLP | $\hat{x}_i[t + s] = \sigma(\mathbf{W}_2 \sigma(\mathbf{W}_1 \mathbf{x}_i^l[t]))$ where $\sigma = \tanh$ is the activation function, $\mathbf{W}_1 \in \mathbb{R}^{H \times l}$, $\mathbf{W}_2 \in \mathbb{R}^{1 \times H}$ are weight matrices, and $\mathbf{x}_i^l[t] = (x_i[t - l + 1], \dots, x_i[t])$. We used $H = 50$ for all patients. |
| Koopman autoencoder<sup>74</sup> | $\hat{x}_i[t + s] = E(\mathbf{x}_i^l[t])^\top \theta + b$ and $\mathbf{x}_i^l[t + 1] = D(KE(\mathbf{x}_i^l[t]))$, where the encoder is $E(\mathbf{x}) = \mathbf{W}_e^2 \sigma(\mathbf{W}_e^1 \mathbf{x})$ with $\mathbf{W}_e^1 \in \mathbb{R}^{p \times l}$, $\mathbf{W}_e^2 \in \mathbb{R}^{p \times p}$, and the decoder is $D(\mathbf{z}) = \mathbf{W}_d^2 \sigma(\mathbf{W}_d^1 \mathbf{z})$ with $\mathbf{W}_d^1 \in \mathbb{R}^{p \times p}$, $\mathbf{W}_d^2 \in \mathbb{R}^{l \times p}$. Here, $\sigma = \text{ReLU}$, $p$ is the latent dimension (set to 64), and $K \in \mathbb{R}^{p \times p}$ is the learnable Koopman operator that evolves the latent state forward by one step. $\theta \in \mathbb{R}^{p \times 1}$ and $b \in \mathbb{R}$. |

### Predictive modeling of seizure risk

The second-level autoregressive (AR) model *ϕ* implements more holistic decision making by looking at historical trends in seizure risk instead of only instantaneous risk. Further, in contrast to simpler techniques such as majority voting or ensemble methods, which aggregate *decisions* from multiple classifiers or time points, the AR model *ϕ* learns sequential trends in data which can lead to more nuanced decision-making. For each patient and baseline model *f* (Eq. (1)), we train *ϕ* as a standard supervised classification problem using the same training data used for learning the baseline model *f* and feature-predictive model *h*. The values of *q*_*B*_[*t k* + 1], …, *q*_*B*_[*t*] are provided as input and the preictal/interictal labels provided as target output. In all cases we have used a random forest classifier for *ϕ* with a lag of *k* = 90 (15 minutes), which we have empirically observed to perform better than alternative methods such as logistic regression and KNN. Once the classifier is trained, for each test sample we obtain *q*_*B*+*R*_[*t*] by computing the ratio of the number of positive (preictal)-classifying tree divided by the number of all decision trees in the random forest.

### Pseudo-prospective analysis

In comparison to the more common approach of evaluating a seizure forecasting model based on pre-segmented, often bal-anced datasets, pseudo-prospective analysis (PPA) measures the accuracy of a seizure forecasting model based on its ability to continuously monitor an iEEG stream over multiple days and correct predict clinically marked seizures while minimizing false alarms during extended interictal periods. Following^72, 75^, we implement pseudo-prospective analysis as follows. Given a test period [*t*_0_, *t*_1_], we first evaluate the seizure risk time series [*q*_*M*_ [*t*_0_], *q*_*M*_ [*t*_0_ + 1], …, *q*_*M*_ [*t*_1_]] where *M* = *B, B*+*F, B*+*R*, or *B*+*F* +*R*. For *M* = *B* and *B* + *R*, this time series is directly shifted by *s* steps to compensate for the lack of an *s*-step ahead predictive component. We then sweep over the value a threshold *τ* between 0 and 1, and for each fixed value of *τ* we calculate the values of sensitivity (true positive rate) and time in warning as follows. At any time *t* ∈ [*t*_0_, *t*_1_], a warning period is triggered when *q*_*M*_ [*t*] exceeds *τ*, and the warning remains active for a fixed duration of 30 minutes following the threshold crossing. This blocking is important in PPA for the stability of the warning raised by the algorithm, as an otherwise “flickering” warning would have little utility and/or lead to major confusions in decision making. The sensitivity is then computed as the fraction of seizures that occur during an active warning period, and time in warning as the percentage of total test time spent in warning. These measures, both a function of *τ*, capture the burdens of false negatives and false positives, respectively.

By varying the threshold *τ* we then obtain the pseudo-prospective sensitivity-time in warning curve, as exemplified in Figure 8. The bottom left and top right corners of this curve are achieved trivially by all models, corresponding to *τ* = 1 and *τ* = 0, respectively. We measure the accuracy of each seizure forecaster based on the area under this curve, denoted by pseudo-prospective area under the curve or PP-AUC. PP-AUC quantifies the trade-off between sensitivity and TIW, allowing for an analysis of how well the model balances early detection and false alarms. PP-AUC ≃ 1 denotes a near-perfect model that correctly predicts all seizures with no false positives, PP-AUC ≃ 0 denotes the opposite, and PP-AUC ≃ 0.3 ∼ 0.4 denotes an approximately at-chance model (chance is often lower than 0.5 because of the temporal blocking in PPA).

### Computing chance-level PP-AUC

Unlike standard AUC for which the chance level is 0.5, even with imbalanced data, chance-level PP-AUC is often less than 0.5 because of the temporal (here, 30min) blocking in PPA. Further, we cannot simply shuffle the test samples to obtain chance level, as seizures typically cluster in time and shuffling would destroy their clustering which is beneficial for any (chance included) seizure forecaster. Therefore, to obtain chance-level PP-AUC we randomize the baseline model *f* by training it over a shuffled training set in which the preictal/interictal labels are randomly shuffled. This randomized model is then tested in PPA as usual, ensuring a fair chance level for PP-AUC that truly reflects a randomized decision maker without altering the structure of PPA.

### Statistical comparison

To allow for statistical comparisons between models, we repeated the classi-fier training multiple times, each time using the preictal periods from the first three lead seizures together with three neighboring interictal segments, and additionally including one or two randomly selected lead seizures (with matched interictal segments) to augment the training set. The corresponding segments were excluded from evaluation, and the remaining iEEG data after the third lead seizure were used for testing. For each random draw of training segments, seizure likelihood estimates from the trained classifier were used to calculate the corresponding PP-AUCs.

## Acknowledgements

The research performed in this study was supported in part by the National Science Foundation Award Number 2239654 (to E.N.) and the National Institute of Health Award Number R01-NS-116504 (to K.A.D.).

## Author Contributions

E.N. designed and supervised the study; G.A. performed the analyses; E.C. performed manual seizure annotations for all patients; K.A.D and E.C. co-supervised the selection and processing of the data and the empirical interpretation of the results; all authors contributed to writing the manuscript.

## Additional Information

The iEEG data used in this study is publicly available from the iEEG.org Portal at https://www.ieeg.org/.

---

**Supplementary Figure 1.**
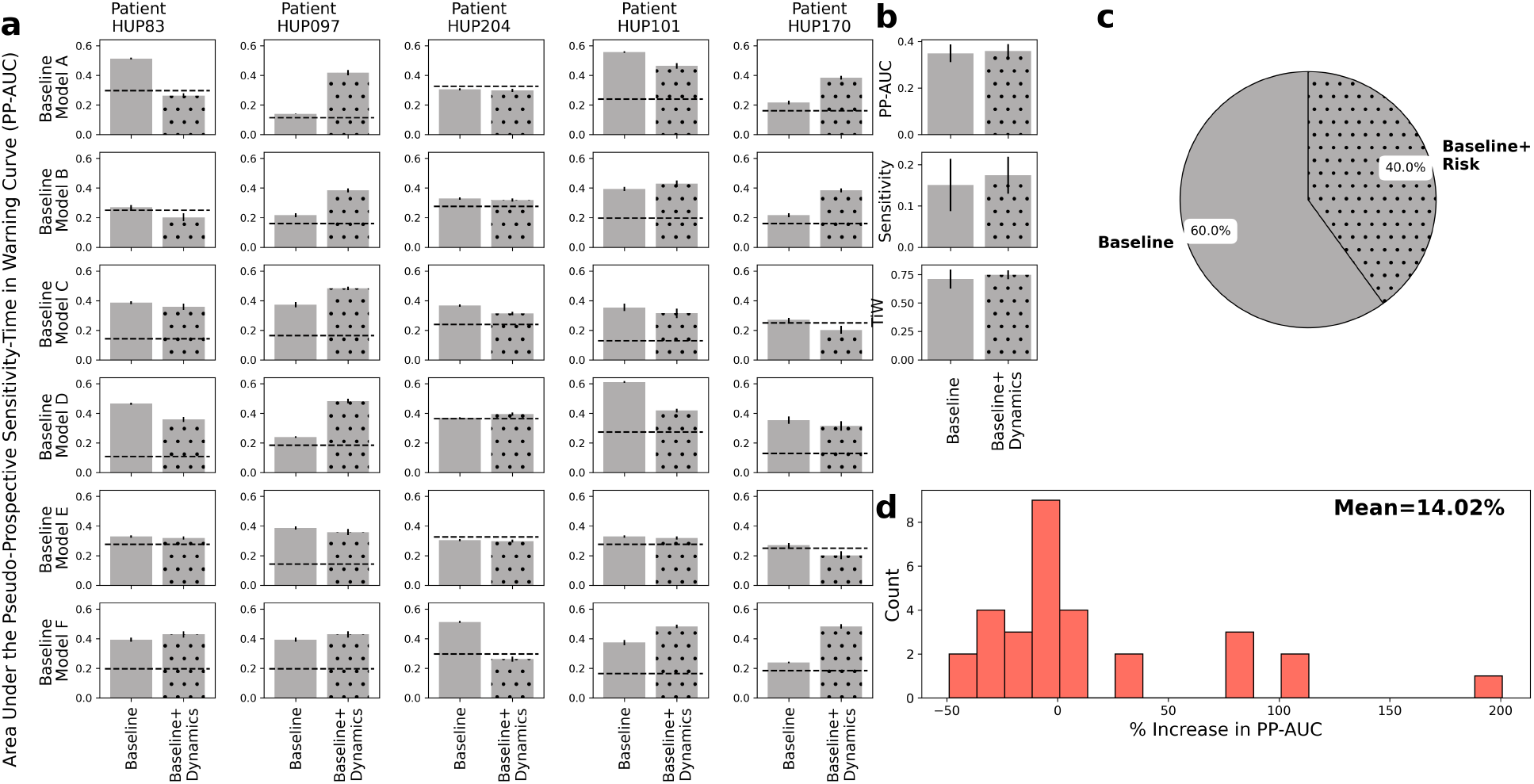
Relative improvements in pseudo-prospective forecasting accuracy using risk dynamical modeling alone. Panels parallel those in Figures 5 and 7. Note the lack of color coding (corresponding to different feature dynamical models which are absent here) and the use of dotted hatches for risk-augmented models.

## Footnotes

† The autoregressive model *ϕ*, while important, is secondary and plays a filtering (smoothing) role.

## References

[1] M.-C. Picot, M. Baldy-Moulinier, J.-P. Daures, P. Dujols, and A. Crespel, “The prevalence of epilepsy and pharmacoresistant epilepsy in adults: a population-based study in a western european country,” Epilepsia, vol. 49, no. 7, pp. 1230–1238, 2008.

[2] A. Fattorusso, S. Matricardi, E. Mencaroni, G. B. Dell’Isola, G. Di Cara, P. Striano, and A. Verrotti, “The pharmacoresistant epilepsy: an overview on existent and new emerging therapies,” Frontiers in neurology, vol. 12, p. 674483, 2021.

[3] A. T. Berg, “Identification of pharmacoresistant epilepsy,” Neurologic clinics, vol. 27, no. 4, pp. 1003–1013, 2009.

[4] B. Litt, R. Esteller, J. Echauz, M. D’Alessandro, R. Shor, T. Henry, P. Pennell, C. Epstein, R. Bakay, M. Dichter et al., “Epileptic seizures may begin hours in advance of clinical onset: a report of five patients,” Neuron, vol. 30, no. 1, pp. 51–64, 2001.

[5] M. O. Baud, T. Proix, N. M. Gregg, B. H. Brinkmann, E. S. Nurse, M. J. Cook, and P. J. Karoly, “Seizure forecasting: bifurcations in the long and winding road,” Epilepsia, vol. 64, pp. S78–S98, 2023.

[6] R. G. Andrzejak, H. P. Zaveri, A. Schulze-Bonhage, M. G. Leguia, W. C. Stacey, M. P. Richardson, L. Kuhlmann, and K. Lehnertz, “Seizure forecasting: Where do we stand?” Epilepsia, vol. 64, pp. S62–S71, 2023.

[7] J. C. Cheng and D. M. Goldenholz, “Seizure prediction and forecasting: a scoping review,” Current Opinion in Neurology, vol. 38, no. 2, pp. 135–139, 2025.

[8] A. Shoeibi, N. Ghassemi, M. Khodatars, M. Jafari, S. Hussain, R. Alizadehsani, P. Moridian, A. Khosravi, H. Hosseini-Nejad, M. Rouhani, A. Zare, A. Khadem, S. Nahavandi, A. F. Atiya, and U. R. Acharya, “Epileptic seizure detection using deep learning techniques: A review,” arXiv preprint arXiv:2007.01276, 2020.

[9] T. Kim, P. Nguyen, N. Pham, N. Bui, H. Truong, S. Ha, and T. Vu, “Epileptic seizure detection and experimental treatment: A review,” Frontiers in Neurology, vol. 11, p. 701, 2020.

[10] A. Sharmila and P. Geethanjali, “A review on the pattern detection methods for epilepsy seizure detection from eeg signals,” Biomedical Engineering/Biomedizinische Technik, vol. 64, no. 5, pp. 507–517, 2019.

[11] Y. Paul, “Various epileptic seizure detection techniques using biomedical signals: a review,” Brain informatics, vol. 5, no. 2, p. 6, 2018.

[12] F. S. Leijten, D. T. Consortium, J. van Andel, C. Ungureanu, J. Arends, F. Tan, J. van Dijk, G. Petkov, S. Kalitzin, T. Gutter, A. de Weerd, B. Vledder, R. Thijs, G. van Thiel, K. Roes, W. Hofstra, R. Lazeron, P. Cluitmans, M. Ballieux, and M. de Groot, “Multimodal seizure detection: A review,” Epilepsia, vol. 59, pp. 42–47, 2018.

[13] bbrinkm and W. Cukierski, “American epilepsy society seizure prediction challenge,” https://kaggle.com/competitions/seizure-prediction, 2014, xkaggle.

[14] L. Kuhlmann, LizLopez, M. O’Connell, rudyno5, S. Wang, and W. Cukierski, “Melbourne university aes/mathworks/nih seizure prediction,” https://kaggle.com/competitions/melbourne-university-seizure-prediction, 2016, xkaggle.

[15] L. Kuhlmann, P. Karoly, D. R. Freestone, B. H. Brinkmann, A. Temko, A. Barachant, F. Li, G. Titericz Jr, B. W. Lang, D. Lavery et al., “Epilepsyecosystem. org: crowd-sourcing reproducible seizure prediction with long-term human intracranial eeg,” Brain, vol. 141, no. 9, pp. 2619–2630, 2018.

[16] E. E. S. Laboratory, “Seizure detection challenge (2025),” Epilepsy Benchmarks website, 2025. [Online]. Available: https://epilepsybenchmarks.com/challenge/

[17] R. E. Stirling, M. J. Cook, D. B. Grayden, and P. J. Karoly, “Seizure forecasting and cyclic control of seizures,” Epilepsia, vol. 62, pp. S2–S14, 2021.

[18] G. Costa, C. Teixeira, and M. F. Pinto, “Comparison between epileptic seizure prediction and forecasting based on machine learning,” Scientific Reports, vol. 14, no. 1, p. 5653, 2024.

[19] M. Dümpelmann, “Early seizure detection for closed loop direct neurostimulation devices in epilepsy,” Journal of neural engineering, vol. 16, no. 4, p. 041001, 2019.

[20] H. Kassiri, S. Tonekaboni, M. T. Salam, N. Soltani, K. Abdelhalim, J. L. P. Velazquez, and R. Genov, “Closed-loop neurostimulators: A survey and a seizure-predicting design example for intractable epilepsy treatment,” IEEE transactions on biomedical circuits and systems, vol. 11, no. 5, pp. 1026–1040, 2017.

[21] A. N. Khambhati, E. F. Chang, M. O. Baud, and V. R. Rao, “Hippocampal network activity forecasts epileptic seizures,” Nature Medicine, vol. 30, no. 10, pp. 2787–2790, 2024.

[22] T. L. Skarpaas, B. Jarosiewicz, and M. J. Morrell, “Brain-responsive neurostimulation for epilepsy (rns® system),” Epilepsy Research, vol. 153, pp. 68–70, 2019.

[23] H. Yang, J. Mueller, M. Eberlein, S. Kalousios, G. Leonhardt, J. Duun-Henriksen, T. Kjaer, and R. Tetzlaff, “Seizure forecasting with ultra longterm eeg signals,” Clinical Neurophysiology, vol. 167, pp. 211–220, 2024.

[24] M. Halimeh, M. Jackson, T. Loddenkemper, and C. Meisel, “Training size predictably improves machine learning-based epileptic seizure forecasting from wearables,” Neuroscience Informatics, vol. 5, no. 1, p. 100184, 2025.

[25] S.-F. Liang, Y.-C. Liao, F.-Z. Shaw, D.-W. Chang, C.-P. Young, and H. Chiueh, “Closedloop seizure control on epileptic rat models,” Journal of neural engineering, vol. 8, no. 4, p. 045001, 2011.

[26] A. Ashourvan, S. Pequito, A. N. Khambhati, F. Mikhail, S. N. Baldassano, K. A. Davis, T. H. Lucas, J. M. Vettel, B. Litt, G. J. Pappas, and D. S. Bassett, “Model-based design for seizure control by stimulation,” Journal of Neural Engineering, vol. 17, no. 2, p. 026009, 2020.

[27] D. Rafik and B. Larbi, “Autoregressive modeling based empirical mode decomposition (emd) for epileptic seizures detection using eeg signals.” Traitement du Signal, vol. 36, no. 3, 2019.

[28] M. S. J. Solaija, S. Saleem, K. Khurshid, S. A. Hassan, and A. M. Kamboh, “Dynamic mode decomposition based epileptic seizure detection from scalp eeg,” IEEE Access, vol. 6, pp. 38 683–38 692, 2018.

[29] W.-C. Chang, J. Kudlacek, J. Hlinka, J. Chvojka, M. Hadrava, V. Kumpost, A. D. Powell, R. Janca, M. I. Maturana, P. J. Karoly, D. R. Freestone, M. J. Cook, M. Palus, J. Otahal, J. G. R. Jefferys, and P. Jiruska, “Loss of neuronal network resilience precedes seizures and determines the ictogenic nature of interictal synaptic perturbations,” Nature neuroscience, vol. 21, no. 12, pp. 1742–1752, 2018.

[30] A. Li, S. Inati, K. Zaghloul, and S. Sarma, “Fragility in epileptic networks: the epileptogenic zone,” in 2017 American Control Conference (ACC). IEEE, 2017, pp. 2817–2822.

[31] A. Aarabi and B. He, “Seizure prediction in hippocampal and neocortical epilepsy using a model-based approach,” Clinical Neurophysiology, vol. 125, no. 5, pp. 930–940, 2014.

[32] L. Chisci, A. Mavino, G. Perferi, M. Sciandrone, C. Anile, G. Colicchio, and F. Fuggetta, “Real-time epileptic seizure prediction using ar models and support vector machines,” IEEE Transactions on Biomedical Engineering, vol. 57, no. 5, pp. 1124–1132, 2010.

[33] M. K. Kiymik, A. Subasi, and H. R. Ozcalik, “Neural networks with periodogram and autoregressive spectral analysis methods in detection of epileptic seizure,” Journal of Medical Systems, vol. 28, no. 6, pp. 511–522, 2004.

[34] L. Logesparan, A. J. Casson, and E. Rodriguez-Villegas, “Optimal features for online seizure detection,” Medical & biological engineering & computing, vol. 50, no. 7, pp. 659–669, 2012.

[35] J. Gotman, “Automatic seizure detection: improvements and evaluation,” Electroencephalography and clinical Neurophysiology, vol. 76, no. 4, pp. 317–324, 1990.

[36] R. Esteller, J. Echauz, T. Tcheng, B. Litt, and B. Pless, “Line length: an efficient feature for seizure onset detection,” in 2001 Conference Proceedings of the 23rd Annual International Conference of the IEEE Engineering in Medicine and Biology Society, vol. 2. IEEE, 2001, pp. 1707–1710.

[37] Z. Zhang and K. K. Parhi, “Low-complexity seizure prediction from ieeg/seeg using spectral power and ratios of spectral power,” IEEE transactions on biomedical circuits and systems, vol. 10, no. 3, pp. 693–706, 2015.

[38] R. Hopfengärtner, B. S. Kasper, W. Graf, S. Gollwitzer, G. Kreiselmeyer, H. Stefan, and H. Hamer, “Automatic seizure detection in longterm scalp eeg using an adaptive thresholding technique: a validation study for clinical routine,” Clinical Neurophysiology, vol. 125, no. 7, pp. 1346–1352, 2014.

[39] M. Saab and J. Gotman, “A system to detect the onset of epileptic seizures in scalp eeg,” Clinical Neurophysiology, vol. 116, no. 2, pp. 427–442, 2005.

[40] M. Bandarabadi, J. Rasekhi, C. Teixeira, and A. Dourado, “Sub-band mean phase coherence for automated epileptic seizure detection,” in The International Conference on Health Informatics. Springer, 2014, pp. 319–322.

[41] M. Hills, “Seizure detection using fft, temporal and spectral correlation coefficients, eigenvalues and random forest,” Github, San Fr. CA, USA, Tech. Rep, pp. 1–10, 2014.

[42] F. Bartolomei, P. Chauvel, and F. Wendling, “Epileptogenicity of brain structures in human temporal lobe epilepsy: a quantified study from intracerebral eeg,” Brain, vol. 131, no. 7, pp. 1818–1830, 2008.

[43] F. T. Sun and M. J. Morrell, “The rns system: responsive cortical stimulation for the treatment of refractory partial epilepsy,” Expert review of medical devices, vol. 11, no. 6, pp. 563–572, 2014.

[44] A. Berényi, M. Belluscio, D. Mao, and G. Buzsáki, “Closed-loop control of epilepsy by transcranial electrical stimulation,” Science, vol. 337, no. 6095, pp. 735–737, 2012.

[45] T. Feng, J. Ni, E. Gleichgerrcht, and W. Jin, “Seizureformer: A transformer model for ieabased seizure risk forecasting,” arXiv preprint arXiv:2504.16098, 2025.

[46] E. Gleichgerrcht, M. Dumitru, D. A. Hartmann, B. C. Munsell, R. Kuzniecky, L. Bonilha, and R. Sameni, “Seizure forecasting using machine learning models trained by seizure diaries,” Physiological measurement, vol. 43, no. 12, p. 124003, 2022.

[47] N. D. Truong, A. D. Nguyen, L. Kuhlmann, M. R. Bonyadi, J. Yang, S. Ippolito, and O. Kavehei, “Convolutional neural networks for seizure prediction using intracranial and scalp electroencephalogram,” Neural Networks, vol. 105, pp. 104–111, 2018.

[48] I. Kiral-Kornek, S. Roy, E. Nurse, B. Mashford, P. Karoly, T. Carroll, D. Payne, S. Saha, S. Baldassano, T. O’Brien, D. Grayden, M. Cook,D. Freestone, and S. Harrera, “Epileptic seizure prediction using big data and deep learning: toward a mobile system,” EBioMedicine, vol. 27, pp. 103–111, 2018.

[49] W. Chen, H. Chiueh, T. Chen, C. Ho, C. Jeng, M. Ker, C. Lin, Y. Huang, C. Chou, T. Fan, M. Cheng, Y. Hsin, S. Liang, Y. Wang, F. Shaw, Y. Huang, C. Yang, and C. Wu, “A fully integrated 8-channel closed-loop neural-prosthetic cmos soc for real-time epileptic seizure control,” IEEE Journal of Solid-State Circuits, vol. 49, no. 1, pp. 232–247, 2013.

[50] A. T. Tzallas, M. G. Tsipouras, and D. I. Fotiadis, “Epileptic seizure detection in eegs using time–frequency analysis,” IEEE transactions on information technology in biomedicine, vol. 13, no. 5, pp. 703–710, 2009.

[51] Y.-x. Zheng, J.-m. Zhu, Y. Qi, X.-x. Zheng, and J.-m. Zhang, “An automatic patient-specific seizure onset detection method using intracranial electroencephalography,” Neuromodulation: Technology at the Neural Interface, vol. 18, no. 2, pp. 79–84, 2015.

[52] A. Shoeb, H. Edwards, J. Connolly, B. Bourgeois, S. T. Treves, and J. Guttag, “Patientspecific seizure onset detection,” Epilepsy & Behavior, vol. 5, no. 4, pp. 483–498, 2004.

[53] S. Heller, M. Hügle, I. Nematollahi, F. Manzouri, M. Dümpelmann, A. Schulze-Bonhage, J. Boedecker, and P. Woias, “Hardware implementation of a performance and energy-optimized convolutional neural network for seizure detection,” in 2018 40th Annual International Conference of the IEEE Engineering in Medicine and Biology Society (EMBC). IEEE, 2018, pp. 2268–2271.

[54] F. Manzouri, S. Heller, M. Dümpelmann, P. Woias, and A. Schulze-Bonhage, “A comparison of machine learning classifiers for energy-efficient implementation of seizure detection,” Frontiers in systems neuroscience, vol. 12, p. 43, 2018.

[55] M. Hügle, S. Heller, M. Watter, M. Blum, F. Manzouri, M. Dumpelmann, A. Schulze-Bonhage, P. Woias, and J. Boedecker, “Early seizure detection with an energy-efficient convolutional neural network on an implantable microcontroller,” in 2018 International Joint Conference on Neural Networks (IJCNN). IEEE, 2018, pp. 1–7.

[56] C. Donos, M. Dümpelmann, and A. Schulze-Bonhage, “Early seizure detection algorithm based on intracranial eeg and random forest classification,” International journal of neural systems, vol. 25, no. 05, p. 1550023, 2015.

[57] J. Yan, J. Li, H. Xu, Y. Yu, and T. Xu, “Seizure prediction based on transformer using scalp electroencephalogram,” Applied Sciences, vol. 12, no. 9, p. 4158, 2022.

[58] K. El Houssaini, C. Bernard, and V. K. Jirsa, “The epileptor model: a systematic mathematical analysis linked to the dynamics of seizures, refractory status epilepticus and depolarization block,” Eneuro, 2020.

[59] M. Guirgis, Y. Chinvarun, M. Del Campo, P. L. Carlen, and B. L. Bardakjian, “Defining regions of interest using cross-frequency coupling in extratemporal lobe epilepsy patients,” Journal of neural engineering, vol. 12, no. 2, p. 026011, 2015.

[60] T. Proix, W. Truccolo, M. G. Leguia, D. King-Stephens, V. R. Rao, and M. O. Baud, “Forecasting seizure risk over days,” MedRxiv, p. 19008086, 2019.

[61] P. J. Karoly, D. M. Goldenholz, D. R. Freestone, R. E. Moss, D. B. Grayden, W. H. Theodore, and M. J. Cook, “Circadian and circaseptan rhythms in human epilepsy: a retrospective cohort study,” The Lancet Neurology, vol. 17, no. 11, pp. 977–985, 2018.

[62] R. Rocamora, R. G. Andrzejak, J. Jiménez-Conde, and C. E. Elger, “Sleep modulation of epileptic activity in mesial and neocortical temporal lobe epilepsy: a study with depth and subdural electrodes,” Epilepsy & Behavior, vol. 28, no. 2, pp. 185–190, 2013.

[63] I. M. Cordeiro, N. Von Ellenrieder, N. Zazubovits, F. Dubeau, J. Gotman, and B. Frauscher, “Sleep influences the intracerebral eeg pattern of focal cortical dysplasia,” Epilepsy research, vol. 113, pp. 132–139, 2015.

[64] M. J. Cook, P. J. Karoly, D. R. Freestone, D. Himes, K. Leyde, S. Berkovic, T. O’Brien, D. B. Grayden, and R. Boston, “Human focal seizures are characterized by populations of fixed duration and interval,” Epilepsia, vol. 57, no. 3, pp. 359–368, 2016.

[65] V. Ferastraoaru, A. Schulze-Bonhage, R. B. Lipton, M. Dümpelmann, A. D. Legatt, J. Blumberg, and S. R. Haut, “Termination of seizure clusters is related to the duration of focal seizures,” Epilepsia, vol. 57, no. 6, pp. 889–895, 2016.

[66] E. Nozari, J. Stiso, L. Caciagli, E. J. Cornblath, X. He, M. A. Bertolero, A. S. Mahadevan, G. J. Pappas, and D. S. Bassett, “Is the brain macroscopically linear? a system identification of resting state dynamics,” bioRxiv, 2020. [Online]. Available: https://www.biorxiv.org/content/early/2020/12/22/2020.12.21.423856

[67] Epilepsy Ecosystem, “Leaderboard,” https://www.epilepsyecosystem.org/leaderboard, accessed: 2025-07-18.

[68] B. Litt and J. Echauz, “Prediction of epileptic seizures,” The Lancet Neurology, vol. 1, no. 1, pp. 22–30, 2002.

[69] J. Tveit, H. Aurlien, S. Plis, V. D. Calhoun, W. O. Tatum, D. L. Schomer, V. Arntsen, F. Cox, F. Fahoum, W. B. Gallentine et al., “Automated interpretation of clinical electroencephalograms using artificial intelligence,” JAMA neurology, vol. 80, no. 8, pp. 805–812, 2023.

[70] Z. Xu, B. H. Scheid, E. Conrad, K. A. Davis, T. Ganguly, M. A. Gelfand, J. J. Gugger, X. Jiang, J. J. LaRocque, W. K. Ojemann et al., “Annotating neurophysiologic data at scale with optimized human input,” Journal of Neural Engineering, 2025.

[71] M. J. Cook, T. J. O’Brien, S. F. Berkovic, M. Murphy, A. Morokoff, G. Fabinyi, W. D’Souza, R. Yerra, J. Archer, L. Litewka et al., “Prediction of seizure likelihood with a long-term, implanted seizure advisory system in patients with drug-resistant epilepsy: a first-in-man study,” The Lancet Neurology, vol. 12, no. 6, pp. 563–571, 2013.

[72] P. J. Karoly, H. Ung, D. B. Grayden, L. Kuhlmann, K. Leyde, M. J. Cook, and D. R. Freestone, “The circadian profile of epilepsy improves seizure forecasting,” Brain, vol. 140, no. 8, pp. 2169–2182, 2017.

[73] B. H. Brinkmann, J. Wagenaar, D. Abbot, P. Adkins, S. C. Bosshard, M. Chen, Q. M. Tieng, J. He, F. Muñoz-Almaraz, P. Botella-Rocamora et al., “Crowdsourcing reproducible seizure forecasting in human and canine epilepsy,” Brain, vol. 139, no. 6, pp. 1713–1722, 2016.

[74] N. Takeishi, Y. Kawahara, and T. Yairi, “Learning koopman invariant subspaces for dynamic mode decomposition,” Advances in neural information processing systems, vol. 30, 2017.

[75] D. E. Snyder, J. Echauz, D. B. Grimes, and B. Litt, “The statistics of a practical seizure warning system,” Journal of neural engineering, vol. 5, no. 4, p. 392, 2008.

